# Multi-omic profiling of transcriptome and DNA methylome in single nuclei with molecular partitioning

**DOI:** 10.1101/434845

**Authors:** Chongyuan Luo, Hanqing Liu, Bang-An Wang, Anna Bartlett, Angeline Rivkin, Joseph R. Nery, Joseph R. Ecker

**Affiliations:** Genomic Analysis Laboratory, The Salk Institute for Biological Studies, La Jolla, CA 92037, United States.; Howard Hughes Medical Institute, The Salk Institute for Biological Studies, La Jolla, CA 92037, United States.

## Abstract

Single-cell transcriptomic and epigenomic analyses provide powerful strategies for unbiased determination of cell types in mammalian tissues. Although previous studies have identified cell types using individual molecular signatures, the generation of consensus cell type classification requires the integration of multiple data types. Most existing single-cell techniques can only make one type of molecular measurement. Here we describe single-nucleus methylcytosine and transcriptome sequencing (snmCT-seq), a multi-omic method that requires no physical separation of DNA and RNA molecules. We demonstrated that snmCT-seq profiles generated from single cells or nuclei robustly distinguish human cell types and accurately measures cytosine DNA methylation and gene expression signatures of each cell type.

## Introduction

Single-cell transcriptome, DNA methylation (5mC) and chromatin profiling techniques have been successfully applied for cell-type classification and studies of gene expression and regulatory diversity in complex tissues. The different techniques used for cell-type identification create a challenge for integration across data modalities. For example, mouse cortical neurons have been studied using various single-cell assays that profile RNA, 5mC or chromatin accessibility ^1–4^, with each study reporting their own classification of cell types. Although it is possible to correlate the major cortical cell types (e.g. cortical layers) identified by transcriptomic and epigenomic approaches, comparing fine level subtypes defined by each of these different methods often yields ambiguous results^2^. Recently, more sophisticated methods based on Canonical Correlation Analysis (CCA) or mutual nearest neighbors (MMN) have been developed to combat batch effects and these methods can potentially be used for integration of different molecular types ^5,6^. However, it is difficult to validate the results of multi-mode comparison solely based on computational analyses, without having a multi-omic reference with different types of molecular measurements made from the same cell.

Existing method such as scM&T-seq and scMT-seq for joint profiling of transcriptome and 5mC rely on the physical separation of RNA and DNA followed by parallel sequencing library preparation ^7,8^. The generation of separate transcriptome and 5mC sequencing libraries leads to a complex workflow and extra costs. It is also unknown whether these methods can be applied to single nuclei, which contain much less polyadenylated RNA than whole cells. Since the cell membrane is ruptured in improperly frozen tissues, the ability to produce robust transcriptome profiles from single nuclei is critically required for applying a multi-omic assay for cell type classification in human tissue specimens.

Here we introduce **m**ethyl**C**ytosine & **T**ranscriptome (mCT-seq) a method that can jointly capture cytosine DNA methylome (5mC) and transcriptome profiles from single cells/nuclei requiring no physical separation of RNA and DNA. This method has been successfully applied for analyzing single cells (scmCT-seq) or single nuclei(snmCT-seq). We demonstrated robust separation of two human cultured cell types - H1 human embryonic stem cells (hESC) and HEK293T cells using either transcriptomic or 5mC profiles extracted from scmCT-seq or snmCT-seq. Transcriptomic and 5mC signatures of H1 and HEK293 cells can be accurately measured using snmCT-seq and snmCT-seq profiles.

## Results

### Joint analysis of RNA and DNA methylome with molecular partitioning

In mCT-seq, RNA and DNA molecules are molecularly partitioned by incorporating 5’-methyl-dCTP instead of dCTP during reverse transcription of RNA (Fig. 1). We use the well-established Smart-seq and Smart-seq2 reactions for cDNA synthesis and the amplification of full-length cDNA (Fig. 1) ^9,10^. The cDNA amplification reaction (second strand synthesis) also incorporates 5’-methyl-dCTP, generating fully methylated double-stranded cDNA amplicons. Using this strategy, all sequencing reads derived from RNA are completely cytosine methylated and do not show the expected C to U sequence changes that would normal occur during treatment with sodium bisulfite. By contrast, more than 95% of cytosines in mammalian genomic DNA are unmethylated and converted to uracils during bisulfite conversion, being read during sequencing as thymine ^11^. In this way sequencing reads originated from RNA and genomic DNA can be distinguished by parsing read-level 5mC density and distribution. Since CpG dinucleotides are highly methylated in mammalian genomes, we used the read-level non-CG methylation (mCH) to sort reads into RNA or DNA partitions. Specifically, we expect all RNA-derived reads to show a 5mC level greater than 90% and DNA-derived reads to show a 5mC level less than 50%. Using this threshold, only 0.04% of single-cell methylome reads were mis-classified as transcriptomic reads whereas only 0.23% ± 0.03% of single-cell RNA-seq reads were mis-classified as methylome reads. These results suggest that RNA-and DNA-derived snmCT-seq reads can be effective separated.

**Figure 1.**
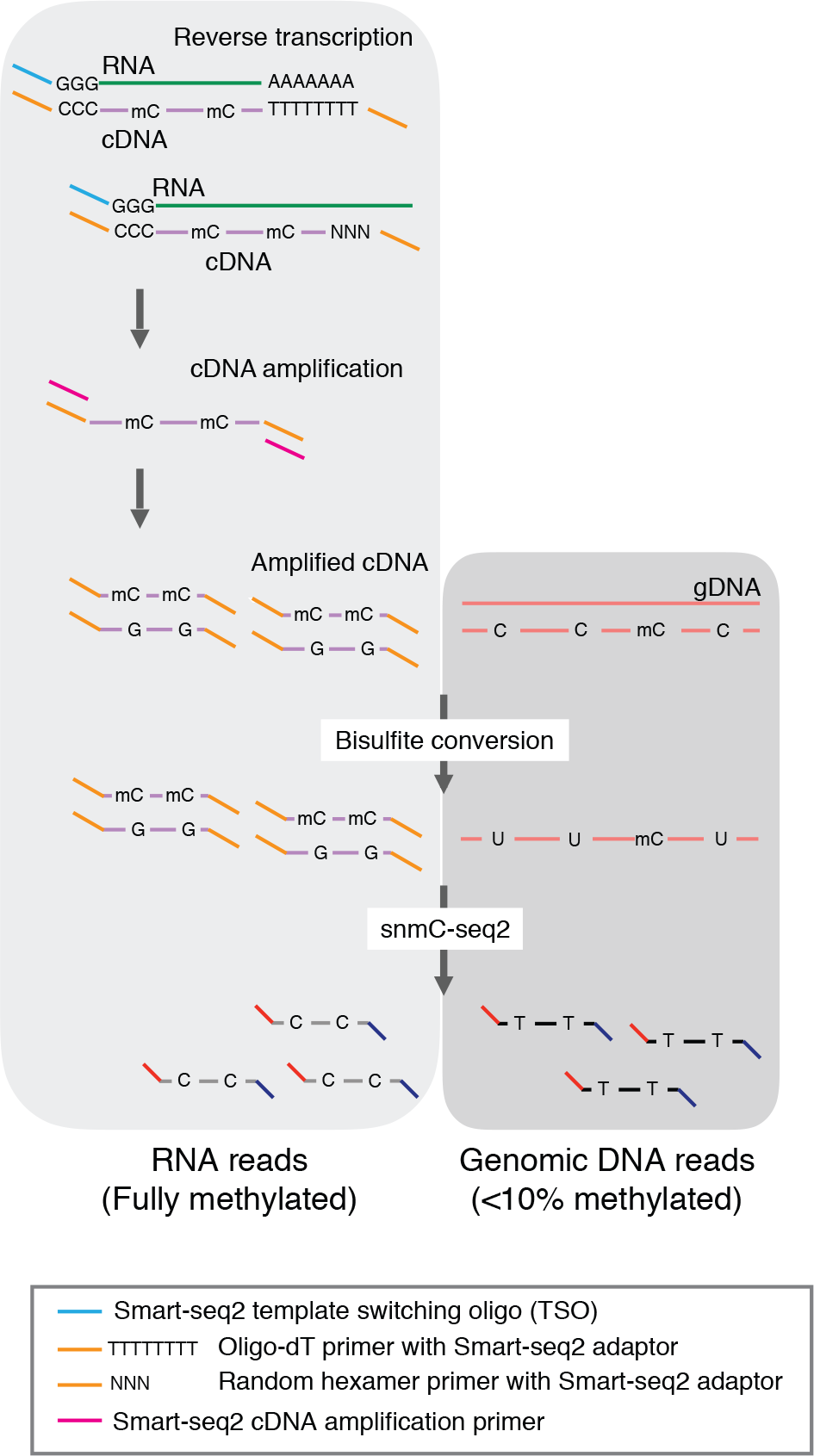
Schematic of mCT-seq shows the molecular partitioning of DNA- and RNA-derived reads.

Following cDNA amplification and bisulfite conversion, sequencing libraries containing both RNA-and DNA-derived molecules are generated using the snmC-seq2 method ^12^. Using a shared library preparation reaction requiring no physical separation of RNA and DNA molecules simplifies the procedure and reduces the cost. In addition, the relative abundance between RNA- and DNA-derived reads can be adjusted by tuning the number of cycles for cDNA amplification.

mCT-seq can be applied to either single whole cells (scmCT-seq) or single nuclei (snmCT-seq). scmCT-seq only uses an oligo-dT primer containing Smart-seq2 adaptor sequence during reverse transcription ^10^, whereas snmCT-seq uses both oligo-dT and random hexamer primers to more efficiently anneal to primary transcripts that often contain introns.

### scmCT-seq and snmCT-seq distinguish human cell types using transcriptome reads

scmCT-seq profiles were generated from 62 H1 hESC and 96 HEK293T cells using 12 cycles of cDNA amplification. H1 and HEK293T cells behaved similarly for both scmCT-seq and snmCT-seq (Supplementary Table 1-2). scmCT-seq detected 4,220 ± 1,251 genes from single whole cells (Fig. 2a). Since nuclei contain much less RNA than whole cells, 15 cycles of cDNA amplification were performed for snmCT-seq. We generated 334 snmCT-seq profiles from an equal mixture of H1 and HEK293 nuclei. H1 and HEK293 nuclei were identified using unbiased clustering and validated using overall 5mC levels and signature genes (e.g. NANOG, LIN28A) specifically expressed in hESC. If only reads mapped to exons were considered, snmCT-seq detected 1,396 ± 863 genes from single nuclei (Fig. 2a). Considering reads mapped to both exons and introns, 4,531 ± 1,888 genes can be detected by snmCT-seq (Fig. 2a). As expected, 17.3 ± 6.1% of snmCT-seq reads were mapped to exons, compared to 68.1 ± 15.2% of scmCT-seq reads mapping to exons (Fig. 2b). Transcriptome reads accounted for 22.2 ± 13.6% and 9.2% ± 6.5% of all mapped reads for scmCT-seq and snmCT-seq, respectively (Fig. 2c). Using a dimension reduction technique t-Distributed Stochastic Neighbor Embedding (t-SNE)^13^, the two human cell types H1 and HEK293T can be readily separated by transcriptomic signatures measured using either scmCT-seq or snmCT-seq (Fig. 1d-e). Further, scmCT-seq or snmCT-seq profiles recapitulate H1 or HEK293 specific gene expression signatures (Fig. 1f).

**Figure 2.**
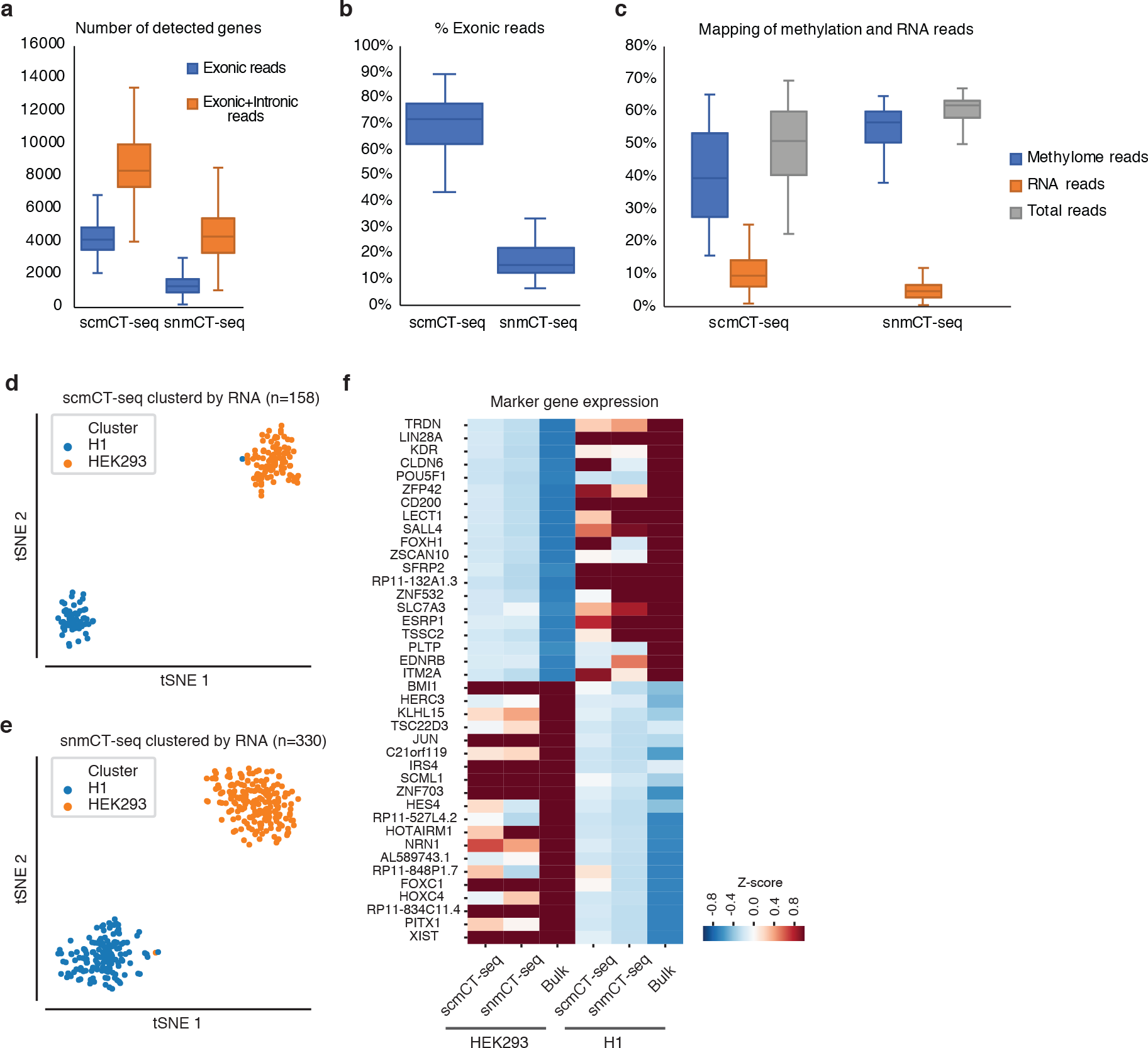
Joint profiling of the transcriptome and DNA methylome detects cell-type specific gene expression signatures. **(a-c)** Number of detected genes (a), percentage of mapped reads that located in exons (b) and mapping rates of methylation and RNA reads (c) for scmCT-seq and snmCT-seq. **(d-e)** Separation of H1 and HEK293T cells by tSNE using transcriptome reads extracted from scmCT-seq (d) or snmCT-seq (e) datasets. **(f)** scmCT-seq and snmCT-seq detect genes specifically expressed in H1 or HEK293T cells.

### scmCT-seq and snmCT-seq distinguish human cell types using DNA methylome reads

To test whether H1 and HEK293T cells can be distinguished using 5mC signatures, tSNE was performed using the average CG methylation (mCG) level of 100 kb non-overlapping genomic bins measured from single cells or nuclei. The two cell types can be readily separated using 5mC profiles extracted from scmCT-seq and snmCT-seq data (Fig. 3a-b). The two single-cell multi-omic assays produced 5mC profiles that are highly consistent to that generated by bulk methylome, as shown by the browser views of pluripotent gene NANOG, and CRNDE locus with enriched expression in colorectal cancer cells (Fig. 3c) ^14^. H1 cells are more heavily methylated (83.6%) in CG context than HEK293T cells (60.1%) as determined from bulk methylomes. In addition, a significant level of mCH (1.3%) is only found in H1 cells but not in HEK293T cells^11^. scmCT-seq and snmCT-seq correctly identified 5mC differences between H1 and HEK293T cells in both CG and non-CG contexts (Fig. 3d-g). To examine whether local 5mC signatures can be recapitulated in scmCT-seq and snmCT-seq profiles, we identified differentially methylated (DMRs) from bulk H1 and HEK293 methylomes. Plotting mCG levels measured using scmCT-seq and snmCT-seq across DMRs showed highly consistent patterns compared to bulk cell methylomes (Fig. 3h-i).

**Figure 3.**
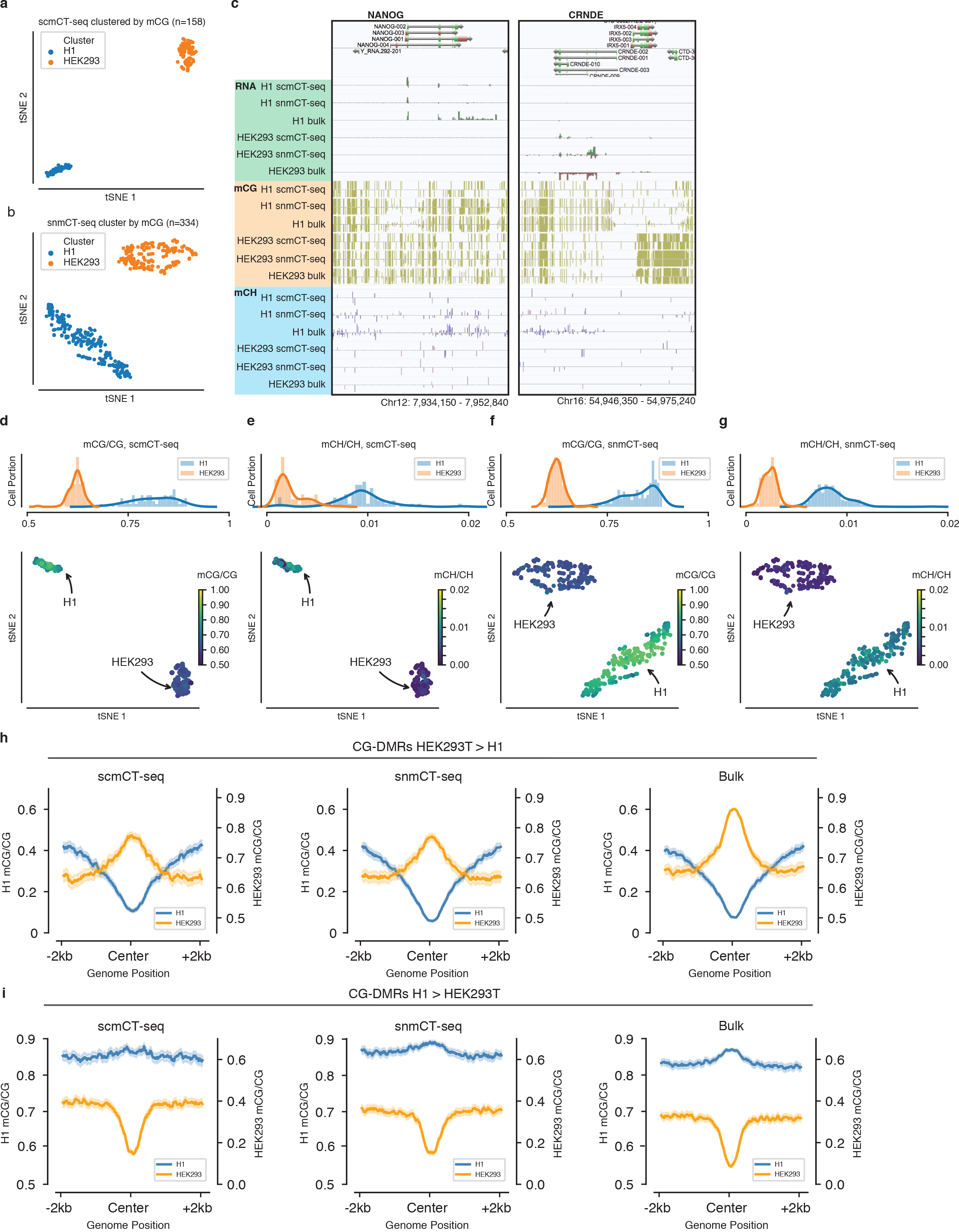
Joint profiling of transcriptome and DNA methylome detects cell-type specific DNA methylation signatures. **(a-b)** Separation of H1 and HEK293T cells by tSNE using DNA methylation information extracted from scmCT-seq (a) or snmCT-seq (b) datasets. **(c)** Browser view of NANOG and CRNDE loci. **(d-g)** Distribution of mCG and mCH levels for single H1 and HEK293 cells/nuclei. **(h-i)** scmCT-seq and snmCT-seq recapitulate bulk mCG patterns at CG-DMRs showing greater mCG levels in HEK293T (h) or H1 (i) cells.

## Discussion

Epigenomic studies often incorporate multiple molecular profiles from the same sample to explore possible correlations between gene regulatory elements and expression. The need for multi-omic comparison poses a challenge for single-cell analysis, since most existing single-cell techniques terminally consume the cell, precluding multi-dimensional analysis. To address this challenge, we have developed a single-cell multi-omic assay snmCT-seq to jointly profile the transcriptome and DNA methylome and can be applied to either single cells or nuclei. snmCT-seq requires no physical separation of DNA and RNA and is designed to be a “single-tube” reaction for steps before bisulfite conversion to minimize material loss. snmCT-seq is fully compatible with high-throughput single-cell methylome techniques such as snmC-seq2^12^ and can be readily scaled to analyze thousands of cells/nuclei.

## Methods

### Cell culture

HEK293T cells were cultured in DMEM with 15% FBS and 1% Penicillin-Streptomycin and dissociated with 1X TrypLE. H1 human ESCs (WA01, WiCell Research Institute) were maintained in feeder-free mTesR1 medium (Stemcell Technologies). hESCs (passage 26) were dispersed with 1U/ml Dispase and collected for single-cell sorting or nuclei isolation. For the sorting of single H1 and HEK293T cells, equal amounts of H1 and HEK293T cells were mixed and stained with anti-TRA-1-60 (Biolegend, Cat#330610) antibody.

### Nuclei isolation from cultured cells

Cell pellets containing 1 million cells were resuspended in 600 μl NIBT [250 mM Sucrose, 10 mM Tris-Cl pH=8, 25 mM KCl, 5mM MgCl_2_, 0.1% Triton X-100, 1mM DTT, 1:100 Proteinase inhibitor (Sigma-Aldrich P8340), 1:1000 SUPERaseIn RNase Inhibitor (ThermoFisher Scientific AM2694), 1:1000 RNaseOUT RNase Inhibitor (ThermoFisher Scientific 10777019)]. The lysate was transferred to a pre-chilled 2 ml dounce homogenizer (Sigma-Aldrich D8938) and dounced using loose and tight pestles for 20 times each. The lysate was then mixed with 400 μl of 50% Iodixanol (Sigma-Aldrich D1556) and gently pipetted on top of 500 μl 25% Iodixanol cushion. Nuclei were pelleted by centrifugation at 10,000 × g at 4°C for 20 min. The pellet was resuspended in 2 ml of DPBS supplemented with 1:1000 SUPERaseIn RNase Inhibitor and 1:1000 RNaseOUT RNase Inhibitor. Hoechst 33342 was added to the sample to a final concentration of 1.25 nM and incubated on ice for 5 min for nuclei staining. Nuclei were pelleted by 1,000 × g at 4°C for 10 min and resuspended in 1 ml of DPBS supplemented with RNase inhibitors.

### Reverse transcription

Single cells or single nuclei were sorted into 384-well PCR plates (ThermoFisher 4483285) containing 1 μl mCT-seq reverse transcription reaction per well. The mCT-seq reverse transcription reaction contained 1X Superscript II First-Strand Buffer, 5mM DTT, 0.1% Triton X-100, 2.5 mM MgCl_2_, 500 μM each of 5’-methyl-dCTP, dATP, dTTP and dGTP, 1.2 μM dT30VN_4 oligo-dT primer (5’-AAGCAGUGGUAUCAACGCAGAGUACUTTTTTUTTTTTUTTTTTUTTTTTUTTTTTVN-3’), 2.4 μM TSO_3 template switching oligo (/5SpC3/AAGCAGUGGUAUCAACGCAGAGUGAAUrGrG+G), 1U RNaseOUT RNase inhibitor, 0.5 U SUPERaseIn RNase inhibitor, 10U Superscript II Reverse Transcriptase. For snmCT-seq, the reaction further included 2 μM N6_2 random primer (/5SpC3/AAGCAGUGGUAUCAACGCAGAGUACNNNNNN). After sorting, the PCR plates were vortexed and centrifuged at 2000 × g. The plates were placed in a thermocycler and incubated using the following program: 25°C for 5 min, 42°C for 90min, 70°C 15min followed by 4°C.

### cDNA amplification

3 μl of mCT-seq cDNA amplification mix was added into each mCT-seq reverse transcription reaction. mCT-seq cDNA amplification reaction contains 1X KAPA 2G Buffer A, 600 nM ISPCR23_2 PCR primer (/5SpC3/AAGCAGUGGUAUCAACGCAGAGU), 0.08U KAPA2G Robust HotStart DNA Polymerase (5 U/μL). A PCR reaction was performed using a thermocycler with the following conditions: 95°C 3min -> [95°C 15 sec -> 60°C 30 sec -> 72°C 2min] -> 72°C 5min -> 4°C. The cycling steps were repeated for 12 to 15 cycles.

### Digestion of unincorporated DNA oligos

1 μl uracil cleavage mix was added to into cDNA amplification reaction. Each 1 μl uracil cleavage mix contains 0.25 μl Uracil DNA Glycosylase (G5010) and 0.25 μl Endonuclease VIII (Y9080) and 0.5 μl Elution Buffer (Qiagen 19086). Unincorporated DNA oligos were digested at 37°C for 30 min using a thermocycler.

### Bisulfite conversion and library preparation

Detailed methods for bisulfite conversion and library preparation are previously described^2,12^. The following modifications were made to accommodate the increased reaction volume of snmCT-seq: Following the digestion of unused DNA oligos, 25 μl instead of 15 μl of CT conversion reagent was added to each well of 384-well plate. 90 μl instead of 80 μl M-binding buffer was added to each well of 384-well DNA binding plate. scmCT-seq libraries were generated using snmC-seq method as described in Luo et al., 2017^2^. snmCT-seq libraries were generated using snmC-seq2 method as described in Luo et al., 2018^12^. The libraries were sequenced using a Illumina HiSeq 4000 instrument with 150 bp paired-end reads.

### Read mapping and the partitioning of transcriptome and methylome reads

To map methylome reads, sequencing reads were mapped to in-silico bisulfite converted hg19 reference genome as previously described ^2^. Mapped reads with MAPQ > 10 were retained for further analyses. Sequencing reads with non-CG methylation level less than 0.5 were considered as true methylome reads. Tab-delimited (allc) files containing methylation level for every cytosine positions was generated using methylpy *call_methylated_sites* function ^15^.

To map transcriptome reads, sequencing reads were mapped to Gencode V19 transcriptome index using STAR 2.5.2b with the following parameters: --alignEndsType EndToEnd --outSAMstrandField intronMotif --outSAMtype BAM Unsorted -- outSAMunmapped Within --outSAMattributes NH HI AS NM MD --sjdbOverhang 100 -- outFilterType BySJout --outFilterMultimapNmax 20 --alignSJoverhangMin 8 -- alignSJDBoverhangMin 1 --outFilterMismatchNmax 999 --outFilterMismatchNoverLmax 0.04 --alignIntronMin 20 --alignIntronMax 1000000 --alignMatesGapMax 1000000 -- outSAMattrRGline ID:4 PL:Illumina. Mapped reads with MAPQ > 10 were retained for further analyses. Non-CG methylation levels for each STAR mapped read was determined from the MD tag. Mapped reads with non-CG methylation levels greater than 0.9 were considered as true transcriptome reads. Transcriptome reads were counted across gene annotations using htseq 0.10.0 with the following parameters: -s no -a 10 -i gene_id -m union. Gene expression was quantified using either only exonic reads with -t exon, or both exonic and intronic reads with -t gene.

The stringency of read partitioning was determined by applying the criteria for identifying snmCT-seq transcriptome reads to snmC-seq2 data (SRR6911760, SRR6911772, SRR6911776) ^12^, which contains no transcriptomic reads. Similarly, the criteria for identifying snmCT-seq methylome reads was applied to Smart-seq data (SRR944317, SRR944318, SRR944319, SRR944320) ^10^, which contains no methylome reads.

### Quantification of DNA methylation

For both snmCT-seq and scmCT-seq datasets, after allc files were generated, the methylated and unmethylated cytosine base calls for cytosines in the CG context were counted for each 100kb bin across genome. All bins with less than 10 total cytosine base calls in CG context were marked as NA. Bins with ＞ 10% of NA were removed. mCG/CG levels were then calculated for each 100kb bin and further normalized by the genome-wide mCG/CG level for each cell. All of the NA values were then replaced with imputed values that are equal to the mean bin value across all cells. To select for highly variable bins and remove outliers, bins with an average mCG/CG > 0.3 and normalized dispersion > 0.2 were retained for further analyses. The normalized dispersion was calculated using SCANPY’s filter_genes_dispersion function ^16^, in which the raw dispersion (variance / mean) of a bin was scaled by standard deviation and mean of regions belong to each bin of mean mCG/CG of regions (n_bins=100). The remaining matrix was transformed with log(X+1) and scaled to mean equal to 0 and variance equal to 1 on each region to produce a mCG matrix.

### Quantification of transcripts abundance

For both snmCT and scmCT datasets, the raw read counts for each whole gene body were transformed into Transcripts Per Kilobase Million (TPM). To select highly variable genes and remove outliers, a method similar to methylation bin filtering was used: genes with mean TPM ∈ [0.1, 50] and normalized dispersion ≥ 0.2 were retained for further analyses. The remaining matrix was transformed by log(X+1) and scaled to mean equal to 0 and variance equal to 1 on each gene to produce an RNA matrix.

### Methylation and RNA Data Clustering and Visualization

For both snmCT and scmCT datasets, PCA was used for dimension reduction of the mCG and RNA matrices. Since only two cell types (H1 and HEK293T) need to be separated, only the first 5 PCs from each matrix were selected to construct K-Nearest Neighbor (KNN) graphs (K=25). On each KNN graph for mCG and RNA, the Louvain method was used to determine clusters (r=0.5) ^17,18^. tSNE was used to visualize the 2-dimensional manifold (perplexity=30) on selected PCs. The Louvain method and tSNE were deployed using functions implemented in SCANPY ^16^.

Clusters were annotated by examining the genome-wide methylation levels and marker gene expression. Data acquired from single cells or nuclei were then merged for each cluster for comparisons with bulk methylome and transcriptome data.

### Comparison to bulk H1 and HEK293 Methylome

The bulk HEK293 cell whole genome bisulfite sequencing (WGBS-seq) data was downloaded from Libertini et. al. (GSM1254259) ^19^. The bulk WGBS-seq data of H1 cell was downloaded from Schultz et al (GSE16256) ^15^. Methylpy was used to call CG-DMRs between these two cell lines^15^. DMRs were filtered by DMS (differentially methylated sites) ≥ 5 and methylation level difference ≥ 0.6.

### Bulk H1 and HEK293 RNA Data Analysis

The bulk HEK293 cell RNA-seq data was downloaded from Aktas et. al. (GSE85161) ^20^, the bulk H1 cell RNA-seq data was downloaded from encodeproject.org (ENCLB271KFE, generated by Roadmap Epigenome). Gene count tables and bigwig tracks were generated using GENCODE v19 gene annotation. In addition, genes with Z-score < 0 in all of the merged cluster RNA profiles were filtered out.

### Data Availability

Methylome and transcriptomic profiles generated by scmCT-seq, snmCT-seq and bulk methylome and transcriptome experiments can be visualized at [http://neomorph.salk.edu/Human_cells_snmCT-seq.php].

## References

1. Zeisel, A. et al. Brain structure. Cell types in the mouse cortex and hippocampus revealed by single-cell RNA-seq. Science 347, 1138–1142 (2015).

2. Luo, C. et al. Single-cell methylomes identify neuronal subtypes and regulatory elements in mammalian cortex. Science 357 600–604 (2017).

3. Preissl, S. et al. Single-nucleus analysis of accessible chromatin in developing mouse forebrain reveals cell-type-specific transcriptional regulation. Nat. Neurosci. 21 432–439 (2018).

4. Tasic, B. et al. Adult mouse cortical cell taxonomy revealed by single cell transcriptomics. Nat. Neurosci. 19 335–346 (2016).

5. Butler, A., Hoffman, P., Smibert, P., Papalexi, E. & Satija, R. Integrating single-cell transcriptomic data across different conditions, technologies, and species. Nat. Biotechnol. 36 411–420 (2018).

6. Haghverdi, L., Lun, A. T. L., Morgan, M. D. & Marioni, J. C. Batch effects in single-cell RNA-sequencing data are corrected by matching mutual nearest neighbors. Nat. Biotechnol. 36 421–427 (2018).

7. Angermueller, C. et al. Parallel single-cell sequencing links transcriptional and epigenetic heterogeneity. Nat. Methods 13 229–232 (2016).

8. Hu, Y. et al. Simultaneous profiling of transcriptome and DNA methylome from a single cell. Genome Biol. 17 88 (2016).

9. Ramsköld, D. et al. Full-length mRNA-Seq from single-cell levels of RNA and individual circulating tumor cells. Nat. Biotechnol. 30 777–782 (2012).

10. Picelli, S. et al. Smart-seq2 for sensitive full-length transcriptome profiling in single cells. Nat. Methods 10 1096–1098 (2013).

11. Lister, R. et al. Human DNA methylomes at base resolution show widespread epigenomic differences. Nature 462 315–322 (2009).

12. Luo, C. et al. Robust single-cell DNA methylome profiling with snmC-seq2. Nat. Commun. 9 3824 (2018).

13. Maaten, L. van der & Hinton, G. Visualizing Data using t-SNE. J. Mach. Learn. Res. 9 2579–2605 (2008).

14. Ellis, B. C., Molloy, P. L. & Graham, L. D. CRNDE: A Long Non-Coding RNA Involved in CanceR, Neurobiology, and DEvelopment. Front. Genet. 3 270 (2012).

15. Schultz, M. D. et al. Human body epigenome maps reveal noncanonical DNA methylation variation. Nature 523 212–216 (2015).

16. Wolf, F. A., Angerer, P. & Theis, F. J. SCANPY: large-scale single-cell gene expression data analysis. Genome Biol. 19 15 (2018).

17. Blondel, V. D., Guillaume, J.-L., Lambiotte, R. & Lefebvre, E. Fast unfolding of communities in large networks. J. Stat. Mech. 2008 P10008 (2008).

18. Levine, J. H. et al. Data-Driven Phenotypic Dissection of AML Reveals Progenitor-like Cells that Correlate with Prognosis. Cell 162 184–197 (2015).

19. Libertini, E. et al. Overexpression of the Heterochromatinization Factor BAHD1 in HEK293 Cells Differentially Reshapes the DNA Methylome on Autosomes and X Chromosome. Front. Genet. 6 339 (2015).

20. Aktaş, T. et al. DHX9 suppresses RNA processing defects originating from the Alu invasion of the human genome. Nature 544 115–119 (2017).

